# Rapid adaptation of an invasive species to climate at a local and continental scale revealed by citizen science

**DOI:** 10.64898/2025.12.10.693511

**Authors:** Riccardo Poloni, Mathieu Joron

## Abstract

Rapid evolution and evolutionary rescue are key mechanisms modulating the success of invasive species, allowing to rapidly adapt to new conditions and to recover from a bottleneck after the introduction. Despite being a well-known mechanism overall, examples of rapid evolution from invasive species and on broad geographic ranges are poorly documented. The box tree moth (*Cydalima perspectalis*) is a polymorphic invasive species that recently colonized Europe, but the ecological factors driving the morph frequencies in natural populations are still largely unknown. Here we took advantage of citizen science, that allows for an unprecedented amount of observations across a large spatial scale, and pheromone trapping, for a more targeted sampling, to detect differences in morph ratios for our model species across Europe and Asia. We then fitted linear regressions and generalized mixed models to detect what variables in the invasive environment drive their frequencies. We found that the melanic form of this moth is maintained at intermediate frequencies in the entire range, but that both at the continental scale and at the local scale, its frequency is higher in warmer and drier conditions. This finding supports the hypothesis of a rapid evolution of colour polymorphism in this invasive species, that established the observed cline in roughly 27 generations. Finally, we provide a physiological rationale of the observed cline, using an experimental approach, showing that melanic individuals have a better resistance to desiccation, compatible with their higher frequency in warmer and drier climates.

## Introduction

Rapid evolution involves evolutionary change occurring on the same time-scale as ecological change. The two processes are classically thought to occur at different time scales (Hairston et al., 2005), but there is growing evidence for rapid evolution being more common than previously thought (Messer et al., 2016). Notably, populations under human-driven pressure like endangered populations, as well as invasive species, tend to experience genetic erosion which can affect their evolutionary potential (Bouzat, 2010; Saccheri et al., 1998). Yet, population growth and adaptation after a contraction are often reported (e.g. Bell & Gonzalez, 2011; Stewart et al., 2017; Tellier et al., 2024) as well as a recovery of genetic diversity (Kaňuch et al., 2021). This rapid adaptation may happen, for instance, via evolutionary rescue, i.e due to natural selection on locally adapted traits such as wing morphology (Hill et al., 1999; Huey et al., 2000), diapause regulation (Byrne & Nichols, 1999), anti-predator strategies (Magurran et al., 1992) or coloration (Endler, 1980; Ni et al., 2024). This often leads to a differentiation of invasive populations compared to the native ones, and compared to the funder population, leading to genetic structure and phenotypic differentiation. Genetic drift may also produce a change in invasive populations during the expansion (Hallatschek et al., 2007; Schlichta et al., 2022) for instance via gene surfing (Hallatschek & Nelson, 2008). Such process involves the random increase in frequency of a “surfing” neutral variant at the invasion front of an expanding population, produced by the higher reproductive success of individuals at the edge of the expansion wave (Hallatschek & Nelson, 2008). It is thus important to consider rapid evolution as part of the eco-evolutionary dynamics of species invasions as a factor impacting the ecology of the community in just a few generations.

The textbook example of rapid evolution is the industrial melanism of the peppered moth, *Biston betularia* (Linnaeus, 1758). This species usually has a white coloration peppered with black dots (*f. typica*), but a completely black colour morph (*f. carbonaria*) started being recorded in the mid 1800s, and became increasingly common, almost entirely replacing the *typica* morph in just a few decades (Majerus, 2009). This increase was caused by a human-induced darkening of resting substrates (trees) and a consequential change in relative camouflage benefits in the two morphs, explaining the increase of *f. carbonaria* (Majerus, 2009; Ruxton et al., 2004).

Industrial melanism in the peppered moth is probably the first documented example of contemporary adaptive evolution driven by anthropogenic change, and a compelling example of the strong selective pressures human activities exert on natural populations. Modern examples of human-driven rapid evolutionary change include rapid adaption to climate change (McCulloch & Waters, 2023), hunting-driven morphological changes (Hard et al., 2008), deforestation, modified predator-prey dynamics (Ni et al., 2024) and many others. Those responses can be triggered by a rapid change in local conditions of a resident population, but also by the translocation of individuals out of their range as in the case of non-native species, which implies a change in local conditions for newly established populations. Therefore, the growing number of species introduced to new territories provides a chance to study contemporary evolution in action and the selective forces shaping their diversity. However, tracking those changes can be challenging at a large spatial scale if the trait under selection is not conspicuous enough. Moreover, although phenotypic trait changes are frequently reported, 70% of such changes have unknown genetic basis (Westley (2011). This can lead to confounding selection with phenotypic plasticity, both important drivers of species invasions (Rohner & Moczek, 2020; Westley, 2011).

Here we study how rapid evolutionary responses arise in the context of the co-invasion of two coexisting, genetically determined colour morphs in the box-tree moth, *Cydalima perspectalis* (Walker, 1859). This moth is native to sub-tropical Asia and arrived in Europe in 2007, spreading very rapidly through the continent (Bras et al., 2019). All invasive populations are polymorphic with two main colour morphs occurring together, a common white morph (*f. typica*) and a rarer melanic morph (*f. fusca*) (Fig. 1), found in variable proportions of about 1:5 melanic:white ratio. A third morph (*f. fasciata*) is also present, but is a minor variation on the *typica* pattern. The forces acting on the maintenance of two discrete, genetically-determined colour morphs during the invasion are still obscure. The fast spread of this polymorphic species at a continental scale provides an excellent system to track rapid character evolution through the invasion and dissect the selective forces acting on it.

**Fig. 1.**
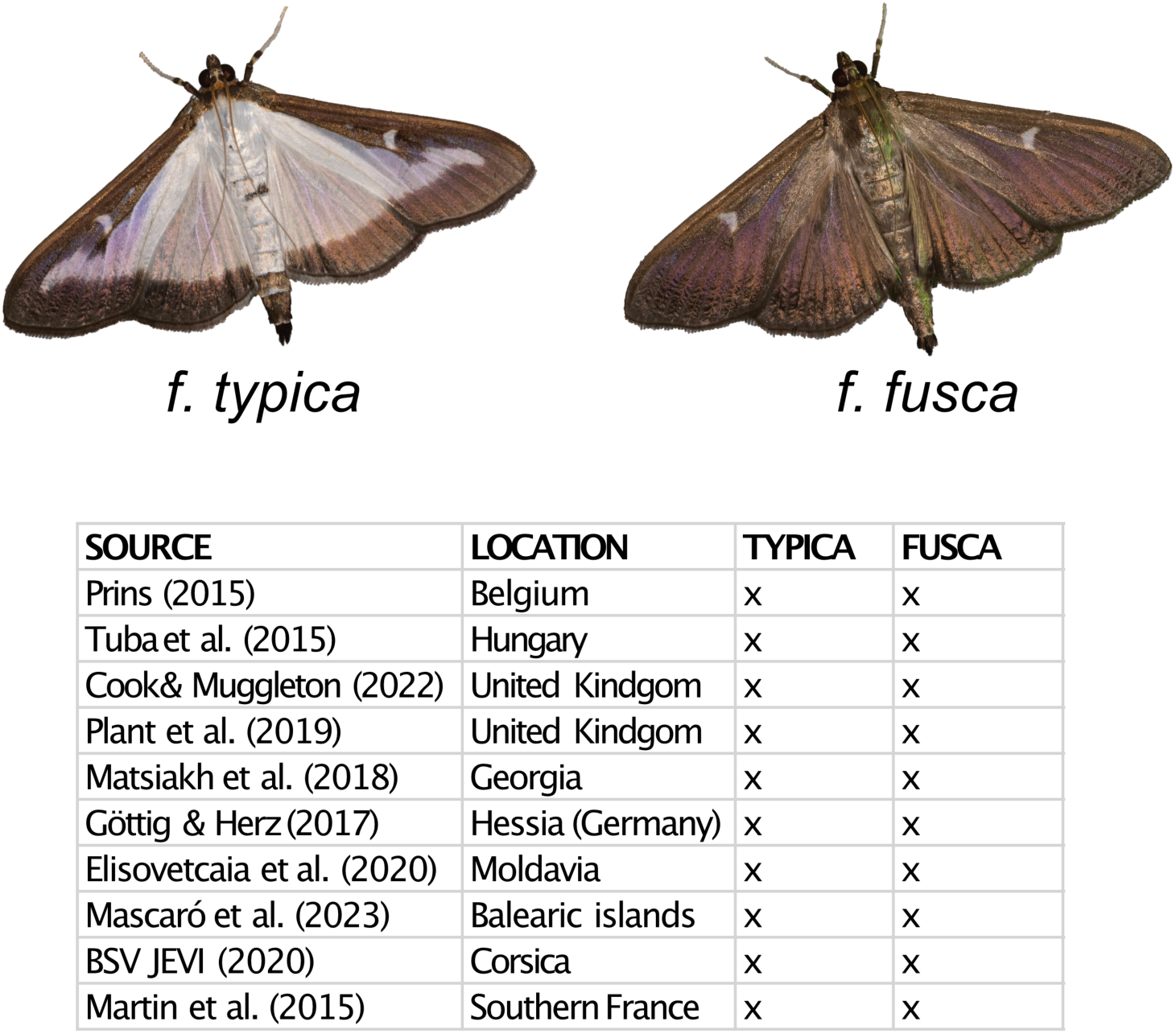
The different colour morphs of the box tree moth (*Cydalima perspectalis*) and the locations in Europe where they have been recorded in the previous literature.

We took advantage of citizen science observations to track the frequency of the two colour morphs of the box tree moth across the continent, allowing, for the first time with citizen science data, not only to track the invasion itself, but also to identify selection on phenotypic traits during the invasion. Using up to ten thousand pictures reported by citizen throughout Europe (invasive range) and Asia (native range). we found that the invasion set up a phenotypic cline associated with certain dimensions of climate variation in Europe. In parallel, standardised morph counts based on pheromone traps in the French Mediterranean revealed similar climatic drive of morph frequencies. Finally, experimental data allowed validating the results obtained by quantifying the influence of climatic factors on the survival of both colour morphs.

## Material and methods

### Morph monitoring using pheromone lures

Pheromone traps were used to determine the variation in morph frequencies in space and time, at a local scale and quantitatively. GinkoBuxus pheromone lures (Sumi Agro, France) were used, which can maintain their efficacy as long as 6 months’ time according to the producer. Traps engineered by INRAE (Institut national de recherche pour l’agriculture, l’alimentation et l’environnement) were used, as they proved more efficient than previous models (Martin et al., 2015). Traps were placed in 12 locations in a 50 km region North from Montpellier (Fig. S1, Table S1), with an altitudinal range of 100-730 m. To mitigate the risk of trap damage or theft, each site received two traps separated by ca. 150 metres, as the pheromone producer suggests spacing traps out by 80-180 metres. Traps were hung from tree branches at 2m above ground, from the end of May to early October, covering the entire flight period of the box tree moth. Each trap was emptied every other week, and samples were preserved frozen until morph counted. White and melanic individuals were counted manually for each site and collection date (Data S1). The same sites were surveyed in 2021 and 2022.

### Citizen science data mining

Occurrences for the box tree moth were obtained from inaturalist (https://www.inaturalist.org/), using only records with photos, for the whole of Europe (invasive range) and from East Asia (native range), which appear to be disjunct ranges. This yielded to 11012 observations for Europe (last accessed, 10^th^ November 2022) and 370 observations for the native range (last accessed, 12^th^ September, 2023; see Table S2 for the specific parameters of the interrogation). This dataset was then filtered to remove coordinates with error margin higher than 1000m, to increase precision when correlating observations with environmental variable values. Each picture was examined individually to check for misidentification and to record morph identity, leading to a curated dataset of 9492 observations for Europe (Fig. S2, Data S2) and 343 for Asia (Data S3). Data for North America, where the box tree moth is also invasive, were not analysed, since most observations are clustered around the city of Toronto, which precludes a comparison between areas with different climatic conditions. Data for Asia were analysed to estimate an indicative morph ratio for the whole native range, but were not sufficient to measure correlation with environmental predictors.

### Desiccation resistance experiments

To test experimentally the hypothesis that morphs might differ in their resistance to desiccation, as suggested by the analysis of citizen science data (see below), we quantified the resistance to desiccation in adults of the box tree moth, belonging to the two morphs. Adults stocks were formed by F1 adults obtained from F0 adult founders bred from caterpillars collected in the same population (Saint-Clément-de-Rivière, France; GPS coordinates 43.718, 3.843) and placed in population cages for mating. F1 caterpillars were fed with fresh *Buxus sempervirens* fresh branches and kept at 20°C. Each pupa, once formed, was removed from the caterpillar cage, weighted with a laboratory precision scale (Pioneer PA413C, 1 mg precision) and placed on a paper tissue in a numbered net-covered, cardboard cup, and placed at 20°C with natural light. Each individual cup was checked twice a day (9 AM, 19:30 PM, and the emergences and deaths were recorded, together with the sex and the morph of the adults. Adults emerging poorly or with visible wing deformations were removed from the experiment (Data S4).

### Statistical analysis

All statistical analysis were performed with R version 2023.03.0 First, to assess whether morph ratios vary in space (between localities) and time (between multiple non-overlapping generations in the same year), data from pheromone traps were analysed, because this quantitative and structured data set allows to extract the effect of phenology and space on morph ratios.

To define successive generations, total number of moths recorded was plotted against collecting dates. The minimum between density peaks was chosen as the boundary between successive generations. To assess morph frequency variations through space and time, a generalized linear model was fitted using morph (white or melanic) as the response variable and locality and generation as explanatory variables, with a binomial distribution and logit link.

Second, to get insight from a broader variation in morph frequencies, climatic conditions, and geographic range, data from citizen science were used. In this case, only changes in morph ratios through space could be tested, because the data did not allow to see how phenology varies in a given place and through the flight period.

Last, to assess whether the two morphs have a different tolerance to desiccation, we fitted a simple generalized linear model using the function glm in lme4 package with lifespan as the response variable and morph, pupal weight and sex as explanatory variable, under a gaussian model. As a second step, since the model retrieved an important contribution of sex, males and females were analysed separately. A Kruskal-Wallis Rank Sum test was used to measure the difference in lifespan between morphs, because female lifespan was not normally distributed.

### Environmental variable correlation

Bioclimatic variables were retrieved from WorldClim as geotiff files (Fick & Hijmans, 2017 - https://www.worldclim.org/ version 2.1) selecting the historical climate dataset, referring to the period 1970-2000, that represents a good approximation of current climate (Fick & Hijmans, 2017) at 30 arc s resolution, corresponding roughly to 1 x 1 km UTM cells. Average UV intensity for the 4 summer months (June, July, August and September, covering most of the moth flight period) was added, obtained by averaging the raster for each month from NASA’s NEO database (https://neo.gsfc.nasa.gov) using the raster calculator tool in QGIS v. 3.10 (A Coruña). A synthetic description of the variables used is given in Table 1.

**Table 1.** Environmental variables used for this study, with their abbreviation and description.

| VARIABLE | DESCRIPTION |
| --- | --- |
| BIO1 | Annual Mean Temperature |
| BIO2 | Mean Diurnal Range (Mean of monthly (max temp - min temp)) |
| BIO3 | Isothermality (BIO2/BIO7) ($\times 100$) |
| BIO4 | Temperature Seasonality (standard deviation $\times 100$) |
| BIO5 | Max Temperature of Warmest Month |
| BIO6 | Min Temperature of Coldest Month |
| BIO7 | Temperature Annual Range (BIO5-BIO6) |
| BIO8 | Mean Temperature of Wettest Quarter |
| BIO9 | Mean Temperature of Driest Quarter |
| BIO10 | Mean Temperature of Warmest Quarter |
| BIO11 | Mean Temperature of Coldest Quarter |
| BIO12 | Annual Precipitation |
| BIO13 | Precipitation of Wettest Month |
| BIO14 | Precipitation of Driest Month |
| BIO15 | Precipitation Seasonality (Coefficient of Variation) |
| BIO16 | Precipitation of Wettest Quarter |
| BIO17 | Precipitation of Driest Quarter |
| BIO18 | Precipitation of Warmest Quarter |
| BIO19 | Precipitation of Coldest Quarter |
| UV_avg_summer | UV index average for summer (june, huly, august, september) |

For each of the 20 bioclimatic variables, the values corresponding to each locality, in the case of pheromone traps, or to each observation, in the case of citizen science data, were extracted. To avoid multicollinearity among variables, two correlation matrices were computed (one for citizen science data and one for pheromone traps, Fig. S3-S4) and variables with a Pearson’s correlation coefficient exceeding 0.85 (Elith et al., 2006) were discarded. Due to a large number of auto-correlated variables, simple generalized linear models were performed on the original set of variables, using morph (white or melanic) as the response variable and one bioclimatic variable at a time. Then, its explanatory power was tested with Chi-Square Anova using Bonferroni correction for multiple testing. However, to avoid removing biologically relevant data (Dormann et al., 2013), those with a known influence on adult moths based on the literature (e.g. Leuthard, 2013; Matošević et al., 2017; Nacambo et al., 2014) were retained regardless of their Pearson’s index and their explanatory power, if they were not redundant (i.e. influencing the same life trait in the study species). Due to the small spatial scale of the pheromone trap dataset, the correlation was very high for most variables (Supplementary Methods, Fig. S4), so we decided to retain the same variables as the citizen science dataset.

With the final set of variables, generalized mixed models were fitted using *glm* function in lme4 package, with morph as response variable and bioclimatic variables as covariates, under a binomial distribution (0 = white morph, 1 = melanic morph) and logit link. The *typica* and *fasciata* morphs were coded as 0 and *fusca* as 1. In the case of citizen-science data, each observation was directly transformed into a binomial variable, and similarly, for the dataset on pheromone traps, each collected moth was considered an observation and then transformed. Four different models were fitted: 1) a model including all variables but not their interactions, 2) a model including the interaction between the variables that are strongly collinear, 3) a model including the interactions among all variables except those with very limited colinearity, and 4) a model including the interactions among all variables. Models were then compared using the Akaike Information Criterion (AIC). For the best model selected, the Hosmer-Lemeshow goodness of fit test was performed which is appropriate for binomial logistic regressions. With the original set of variables linear regressions were also fitted using *lm* function from base R. The frequency of *fusca* individuals was used as an independent variable, and the environmental variables as dependent variables. In the case of pheromone dataset, *fusca* morph frequency was simply obtained by averaging the frequencies recorded by year. In the case of citizen science dataset, since observations were scattered, they didn’t allow to compute a local morph ratio. Therefore, first, environmental variable values were rounded to 0 or 1 decimals, based on how much variance the variable was showing in the dataset, and then the morph ratio per each variable value were computed. This allowed to create variable classes with enough variance to see eventual correlations between morph ratios and climatic variables, but avoiding too accurate measures that would have been associated to just a few observations. Regressions were then evaluated using adjusted R-square value.

The data used in this paper can be found at: https://github.com/r-poloni/rapid_evol The code will be shared upon peer-review publication.

## Results

### Phenology

Data on pheromone traps indicate a bimodal phenology, supporting the existence of two main generations in the study area (Fig. 2). Low elevation sites, however, display a weak increase in captures after a 0 minimum at the end of august 2022, compatible with a partial third generation, with abundances an order of magnitude lower than the June and July peaks (Data S1). A minor phenological advance is observed at low elevation sites compared to the higher elevation sites (Data S1 and Fig. 2).

**Fig. 2.**
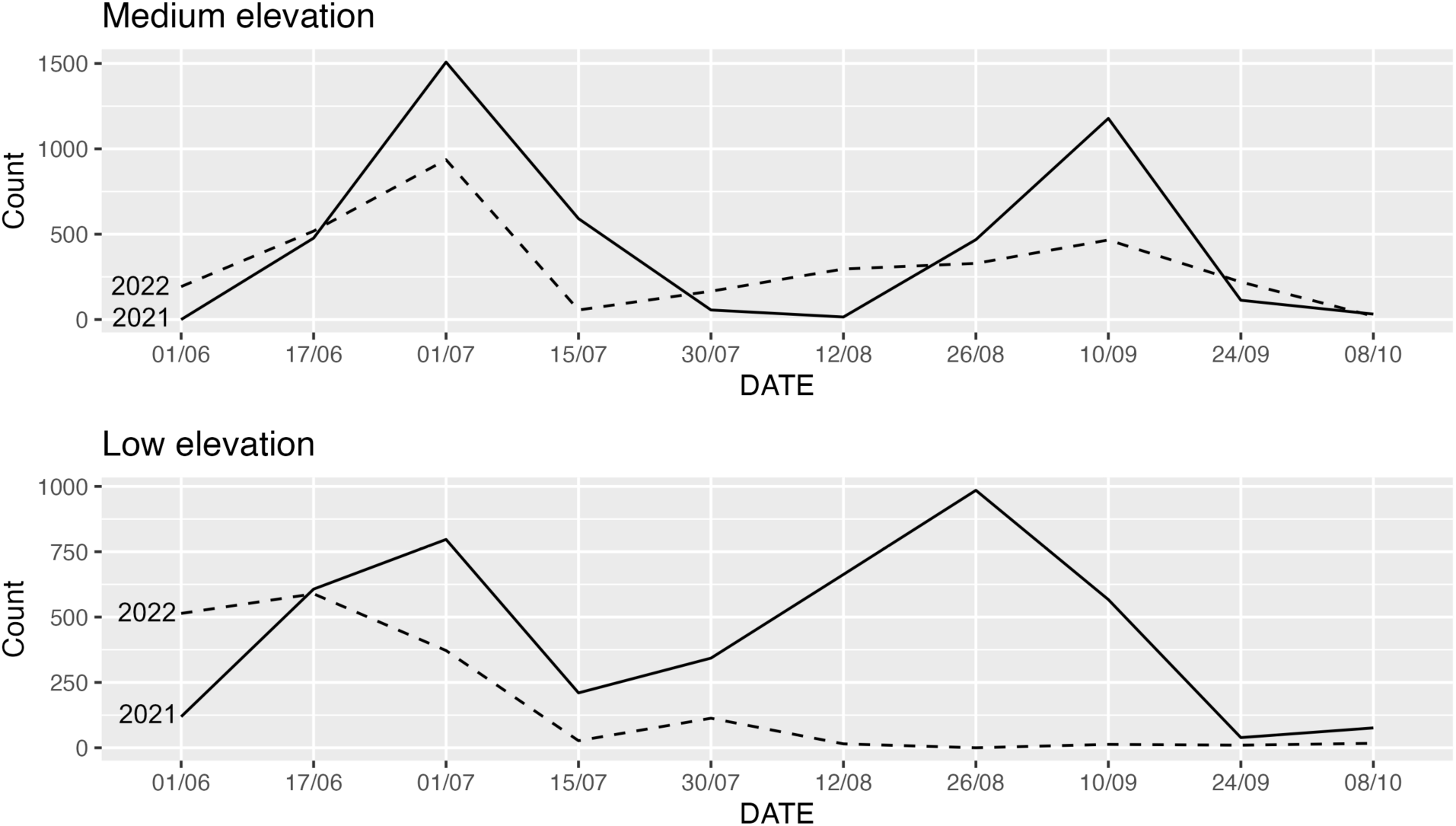
Adult box tree moth phenology through their flight period, as inferred from pheromone traps individual count in 2021 and 2022.

### Morph ratios

The average frequency of the melanic morph in pheromone traps is 0.17 [0.14-0.23]. The average melanic frequency at the European scale, as obtained from citizen science data, is around 0.15, and 0.05 for Asia.

In the GLM, generation did not explain variations in morph ratios (Anova Chi-Square test, p = 0.355), whereas locality explained most of the variance (Anova Chi-Square test, p < 0.001).

### Environmental variables

After variable selection (i.e. preliminary single-variable models and multicollinearity analysis) a total of 7 variables were retained for both datasets, namely annual mean temperature (bio_1), mean diurnal range (bio_2), temperature annual range (bio_7), mean temperature of warmest quarter (bio_10), precipitation seasonality (bio_15), precipitation of warmest quarter (bio_18) and UV intensity for summer months (UV_avg_summer). Following preliminary models, 11 variables were discarded, and another 3 were discarded after multicollinearity analysis.

Among the generalized linear models performed on the citizen science dataset, model 2 was the best (AIC = 6052, Table 2), including interactions between all variables except bio_2 and bio_15 which showed low collinearity with the other variables (Fig. S5). The Hosmer and Lemeshow goodness of fit (GOF) supports a good fit for this model (p = 0.576). The variables with significant effects in the model are bio_1, bio_7, bio_18 and UV_avg_summer as well as the interactions, visible in Table 2. This was expected, because of significant collinearity between them (Fig. S5).

**Table 2.** Chi-Square Anova results for model 2 fitted on the citien science dataset, including interactions between all variables except bio_2 and bio_15. Only variables emerging as significant are shown.

|  | Df | Deviance | Resid. Df | Resid. Dev | Pr(>Chi) |
| --- | --- | --- | --- | --- | --- |
| **bio_1** | 1 | 25.47 | 7871.00 | 6132.33 | < 0.001 |
| **bio_7** | 1 | 16.10 | 7870.00 | 6116.23 | < 0.001 |
| **bio_18** | 1 | 5.78 | 7868.00 | 6110.45 | 0.016 |
| **UV_avg_summer** | 1 | 18.78 | 7867.00 | 6091.67 | < 0.001 |
| **bio_1:bio_10** | 1 | 6.96 | 7863.00 | 6081.50 | 0.008 |
| **bio_7:bio_18** | 1 | 10.00 | 7860.00 | 6070.74 | 0.002 |
| **bio_10:bio_18** | 1 | 15.49 | 7859.00 | 6055.25 | < 0.001 |
| **bio_1:UV_avg_summer** | 1 | 19.34 | 7858.00 | 6035.91 | < 0.001 |
| **bio_10:UV_avg_summer** | 1 | 5.94 | 7856.00 | 6027.28 | 0.015 |
| **bio_18:UV_avg_summer** | 1 | 10.56 | 7855.00 | 6016.72 | 0.001 |
| **bio_1:bio_7:UV_avg_summer** | 1 | 3.87 | 7850.00 | 6009.14 | 0.049 |
| **bio_1:bio_18:UV_avg_summer** | 1 | 6.20 | 7847.00 | 6002.75 | 0.013 |
| **bio_10:bio_18:UV_avg_summer** | 1 | 4.15 | 7845.00 | 5996.68 | 0.042 |
| **bio_1:bio_10:bio_18:UV_avg_summer** | 1 | 8.16 | 7841.00 | 5985.48 | 0.004 |

The best model explaining morph ratios in the pheromone trap dataset was model 1, i.e. with no interactions (AIC = 12228, Table 3). The Hosmer and Lemeshow goodness of fit test supports a good fit for this model (p = 0.884). Only bio_1 and bio_15 significantly explained the probability of detecting a *fusca* individual. The low number of variables with significant effects, compared with the citizen science dataset, may reflect the lower variability of environmental variables, at such a limited spatial scale.

**Table 3.** Chi-Square Anova results for model 1 fitted on the pheromone traps dataset, including no interactions. Only variables emerging as significant are shown.

|  | Df | Deviance | Resid. Df | Resid. Dev | Pr(>Chi) |
| --- | --- | --- | --- | --- | --- |
| **bio_1** | 1 | 25.63 | 13712.00 | 12220.14 | < 0.001 |
| **bio_15** | 1 | 4.17 | 13708.00 | 12213.93 | 0.041 |

Among the linear regressions fitted on the citizen science dataset, variables bio_1, bio_2, bio_5, bio_10, bio_14 and UV_avg_summer emerged as significant (Table 4, Fig. 3). Variable bio_2, however, had a relatively high p-value (p = 0.016) and a very poor correlation coefficient (adjusted R-squared = 0.15). Finally, bio_15 and bio_18 yielded poorly representative plots (Fig. 3). The variable that best explained the melanic morph ratio is bio_1 (i.e. average yearly temperature, adjusted R-square = 0.83, p < 0.001). UV_avg_summer (i.e., the average UV index for June, July, August and September) and bio_5 (i.e. maximum temperature of the warmest month) were also strongly correlated with melanic morph frequency but had an overall lower R-squared value (Table 4). In additionboth bio_5 and UV_avg_summer were strongly collinear with bio_1.

**Fig. 3.**
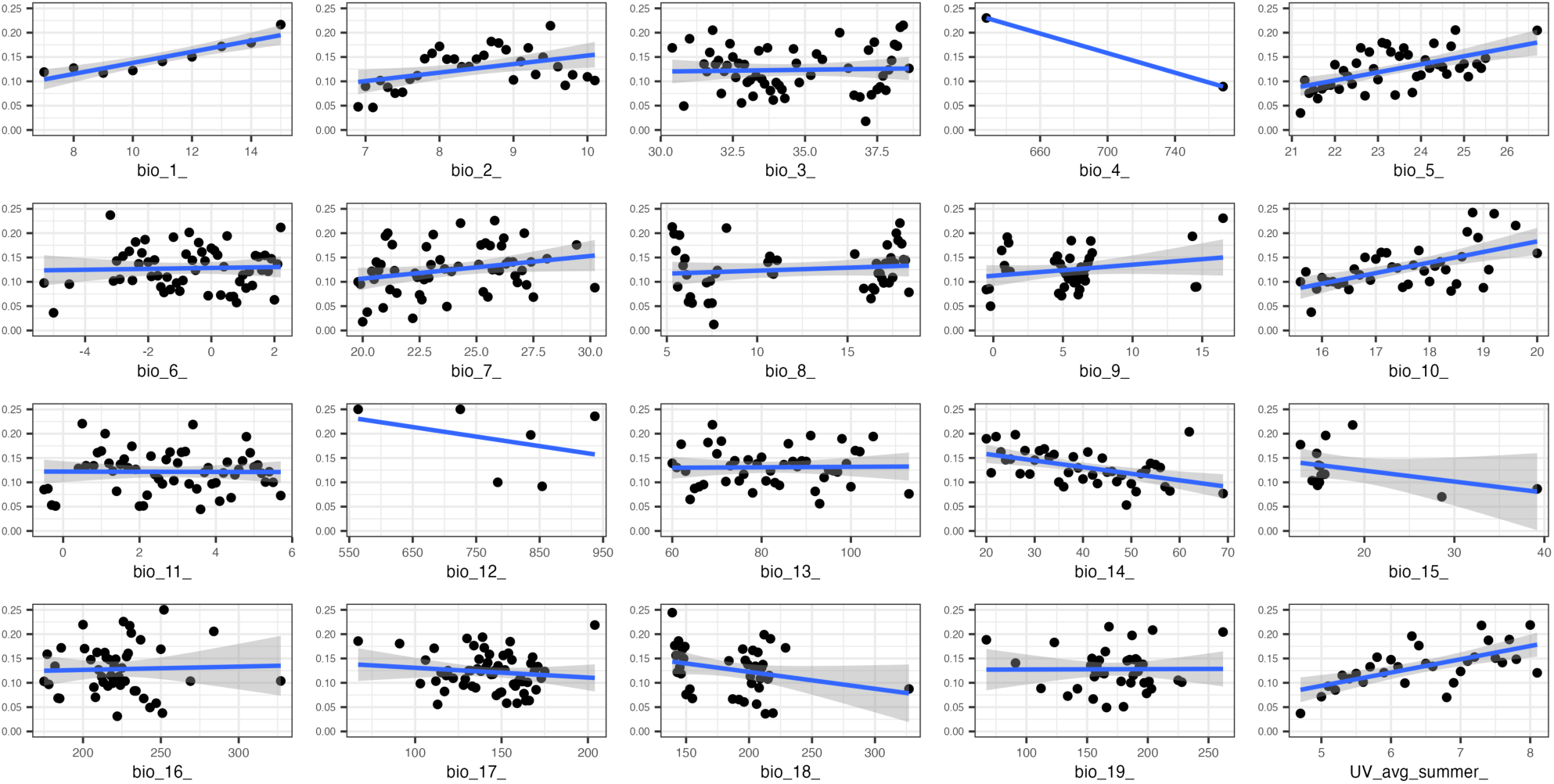
Linear regressions performed using citizen science data. The y axis represents *fusca* morph ratio, the x axis the variable value. The grey shade indicates the confidence intervals of the regression.

**Table 4.**
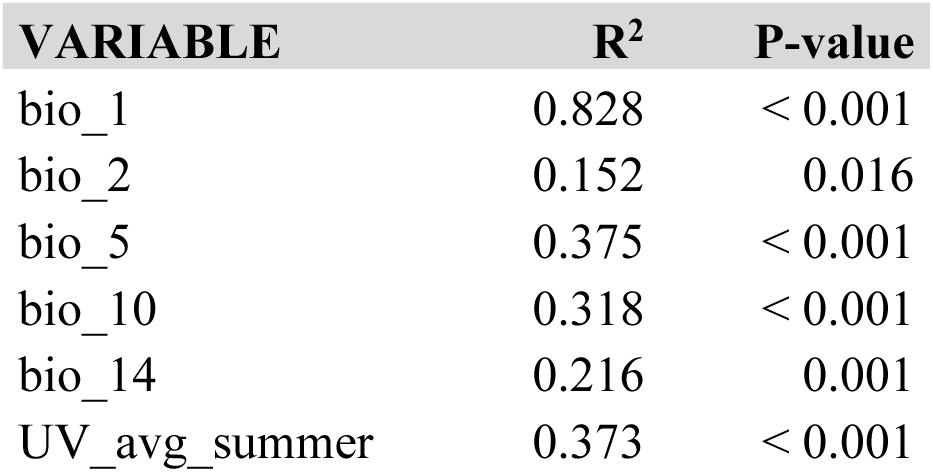
Correlation coefficient (adjusted R square) and p-value for the linear regressions performed on the citizen science dataset. Only variables emerging as significant are shown.

Among the linear regressions fitted on the pheromone trap dataset, all variables except bio_2, bio_3, bio_4 and bio_7 emerged as significant (Table 5, Fig. 4), but bio_7 yielded poorly representative plots and was not considered further here (Fig. 4). The variable that best explained the melanic morph frequency was bio_15, followed by a few other variables all showing good predictive power (Table 5). Unfortunately, disentangling the relative importance of each variable, when comparing different regressions performed with the same dependent variable, is extremely challenging.

**Fig. 4.**
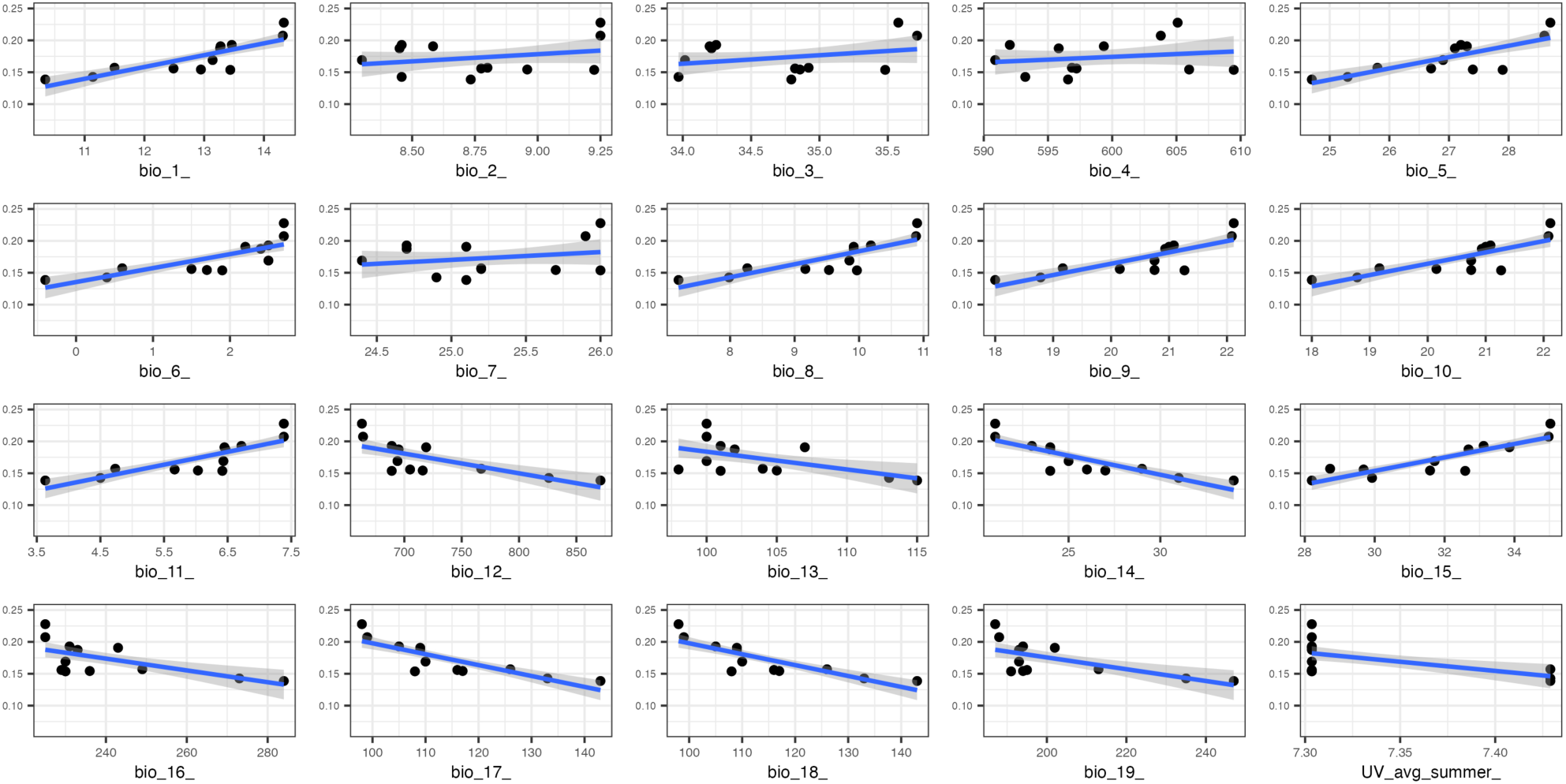
Linear regressions performed using pheromone traps data. The y axis represent fusca morph ratio, the x axis the variable value. The grey shade indicates the confidence intervals of the regression.

**Table 5.** Correlation coefficient (adjusted R square) and p-value for the linear regressions performed on the pheromone traps dataset. Only variables emerging as significant are shown.

| VARIABLE | R<sup>2</sup> | P-value |
| --- | --- | --- |
| bio_1 | 0.661 | < 0.001 |
| bio_5 | 0.589 | < 0.001 |
| bio_6 | 0.628 | < 0.001 |
| bio_8 | 0.685 | < 0.001 |
| bio_9 | 0.643 | < 0.001 |
| bio_10 | 0.643 | < 0.001 |
| bio_11 | 0.683 | < 0.001 |
| bio_12 | 0.495 | < 0.001 |
| bio_13 | 0.259 | 0.007 |
| bio_14 | 0.708 | < 0.001 |
| bio_15 | 0.757 | < 0.001 |
| bio_16 | 0.380 | 0.001 |
| bio_17 | 0.683 | < 0.001 |
| bio_18 | 0.683 | < 0.001 |
| bio_19 | 0.387 | 0.001 |
| UV_avg_summer | 0.314 | 0.003 |

Overall, analyses on both the citizen science and pheromone traps datasets agree on the importance of mean temperature and precipitation, namely showing a positive correlation between *fusca* morph ratio and temperature (bio_1), and a negative correlation with precipitation (bio_14 and bio_15) as visible in Figs. 3-4 and Tables 2-5. Uv index was also positively correlated to the *fusca* ratio in both datasets.

### Desiccation resistance

In the generalized liner model fitted with lifespan as the response variable and morph, pupal weight and sex as explanatory variable both morph and sex emerged as significant effects (Anova Chi-Square test, p = 0.002 for morph and p = 0.003 for sex, Table S3), but not their interaction. In each sex, lifespan was significantly higher in *typica* than in *fusca* morphs (Wilcoxon’s signed-rank test, p = 0.001 for males, p = 0.031 for females), although lifespan variance for the females was much higher in females than in males (mean = 4.95, variance = 1.73 for females, vs mean = 4.13, variance = 0.75 for males, Fig. 5).

**Fig. 5.**
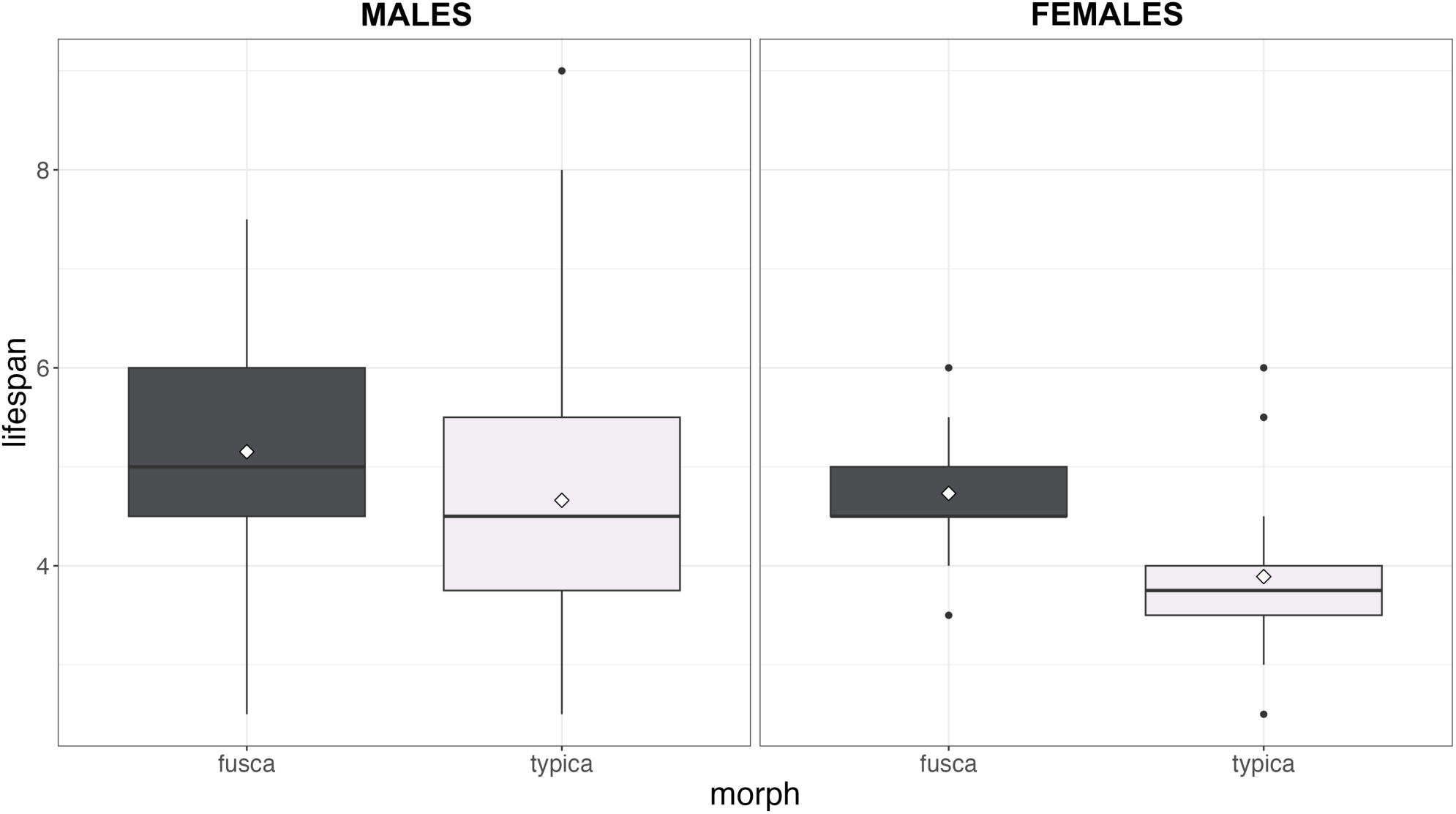
Differences in lifespan (y axis) between the two colour morphs *typica* and *fusca*, each sex analyzed separately. The white diamond in the boxplots represents the mean, the solid line the median.

## Discussion

### Two datasets, two spatial scales

The response of morph ratios of the box tree moth to environmental variables was investigated using two complementary approaches. One approach leveraged opportunistic citizen science observation data that spans a wide spatial scale and the complete array of environmental conditions encountered by the invasion, allowing to explore macro-ecological patterns. A second approach was based on local sampling using pheromone lures, yielding structured data and accurate local morph frequencies, yet with a narrower spatial and ecological scale, allowing to explore local ecological patterns.

Our results showed some differences between the citizen science and the pheromone traps datasets and between GLMs and linear regressions. While some variables were consistently found to contribute to the model, the main differences lied in their relative importance, and some variables were retrieved as significant only in some models. One major source of uncertainty in the case of generalized linear models is multi-collinearity among environmental predictors. Multi-collinearity was expected, because many variables in the WorldClim dataset are correlated (Fick & Hijmans, 2017), and because of the two different spatial scales (Braunisch et al., 2013; Dormann et al., 2013). Removing collinear predictors may remove multi-collinearity, but in the absence of pre-existing data on the species niche, it is considered preferable to leave additional variables that may contribute to the final models (Braunisch et al., 2013). We thus removed strongly collinear variables while retaining weakly collinear variables and those that emerged as highly explicative in the preliminary models or particularly relevant for the species biology. By doing this, however, the power of binomial regressions was reduced when interactions were not considered and so was our ability to untangle the relative contributions of the variables. Multicollinearity in the linear regressions could be partially solved with a PCA regression, but this was only possible for the structured dataset (traps), since the citizen science analysis required grouping observations into polygons with the same variable value, and such polygons were different for every variable.

Despite those analytical differences though, both approaches point in the same direction, and concur in revealing a positive relationship between melanic frequency and temperature and summer UV intensity, and a negative relationship with precipitation. This consensus supports a strong and pervasive effect of climatic drivers on morph ratios in this moth. Melanic alleles are at higher frequency in drier and hotter places, an effect detected at the continental scale in both the invasive and native ranges, as well as when comparing localities within a small part of the invasive range. This analysis therefore reveals the effect, at an ecological scale, of climate on the population genetics of invasive populations.

### Natural selection vs genetic drift

Allelic frequencies may change during an invasion mainly via selection or drift, with contrasting effects. Variants affected by drift via gene surfing usually distribute heterogeneously in the landscape, and with a low frequency in the invasion centre (Hallatschek et al., 2007), which in the case of the box tree moth is Southern Germany. However, morph ratios available from the literature and from our data do not support this (Figs. 1, 4, 5). It is possible though that the higher *fusca* morph frequency in Europe (17%) versus Asia (5%) reflects an initial population bottleneck and the action of random genetic drift at the introduction point (founder effect). However, observed morph ratios within Europe now follow a clear cline set up along an environmental gradient across the invasion range, which contradicts gene surfing. Finally, the *fusca* morph brings adaptive survival benefits in a context of resource depletion, in line with the pattern shown by environmental variables, and cannot be considered neutral as in the case of variants affected by gene surfing (Hallatschek et al., 2007; Hallatschek & Nelson, 2008). Overall, the evidence points for an influence of selection on the morph ratios in the invasive range.

This evolutionary change was extremely rapid: considering an introduction in 2007, a colonization of nearly all the European territory in 2016 (Bras et al., 2019), and an average of 2.5 generations per year (2 to 3 generations/year depending on area), we can estimate that the observed morph ratios evolved in 27 ± 9 generations (Göttig & Herz, 2017; Nacambo et al., 2014). Such rapid change may reflect strong selective pressure, including some related to climatic conditions, as reported for other contemporary evolution models (Catullo et al., 2019). According to Nacambo et al. (2014), the European invasive territory is climatically less suitable to the box tree moth than its native range, and this species is invasive in territories with overall low predicted suitability. This may suggest that the box tree moth settled out of its ecological optimum, perhaps aided by a lack of competitors and predators (e.g. parasitoids), but also owing to rapid adaptation to local climatic conditions, as reported in a similar invasion (Dai et al., 2023).

### Selective pressures acting on the polymorphism

Designed to test experimentally the hypothesis that melanic morph obtain a selective advantage in hot and dry climates, our experiments indeed revealed that the melanic morph *fusca* displayed a better resistance to desiccation compared with the white morph *typica*. The larger variance in lifespan displayed by females compared to males may reflect behavioural variation, for instance in the intensity of flight activity. Overall, melanic individuals are fitter in conditions set to mimic drought periods in climates with dry summers. This corresponds to a thermal melanism hypothesis, since melanisation may associate with more sclerotized cuticle and a better resistance to dehydration (Kalmus, 1941; Välimäki et al., 2015).

In addition, the *fusca* morph lacks the UV reflection found on the wings of the *typica* morph, and is less attacked by predators due to its lower conspicuousness (Poloni et al., 2024). Moreover, studies suggest that UV reflectance may contribute to enhanced conspicuousness of butterfly wings (e.g. Chan et al., 2019). This may lead to an increased conspicuousness of the *typica* morph, that is the more conspicuous already, in areas with high UV reflectance, with a consequent higher predation pressure. This hypothesis is compatible with our results, showing a higher proportion of the *fusca* morph in areas with a higher UV reflectance.

In conclusion, our study reveals a fascinating example of how climatic drivers may drive a rapid evolutionary change and adaptation to new environmental conditions at a continental and local scale. Such adaptation may have increased the invasive potential of the box tree moth in Europe, which we revealed using an original combination of opportunistic citizen science data, standardized population survey, and laboratory experimental tests. Finally, our analysis demonstrates that citizen science can be a reliable and easily accessible source of distributional data, that can be used to track invasive routes and detect evolutionary change occurring during invasions.

## Author Contributions

**Riccardo Poloni**: conceptualization, methodology, field work, investigation, data curation and formal analysis, writing; **Mathieu Joron**: conceptualization, funding acquisition, methodology, field work, investigation, writing.

## Acknowledgements

We are deeply indebted to Elisabeth Tabone, Emeline Morel and Laurent Pelozuelo, for the suggestions about box tree moth pheromones and monitoring. We thank Lola George, Paul Doniol-Valcroze, and the experimental facility at CEFE, for help during the experiments. We warmly thank Yann Le Poul, Paul Cuchot, Allowen Evin and Pierre Lacoste for comments about the analyses, and especially to Francesco Cerasoli, for his advice about the statistical framework of the paper. We are thankful to Robert Wilson and Roger Vila for their useful comments about the first results presented in this paper.

