## Supplementary material for "Rapid adaptation of an invasive species to climate at a local and continental scale revealed by citizen science"

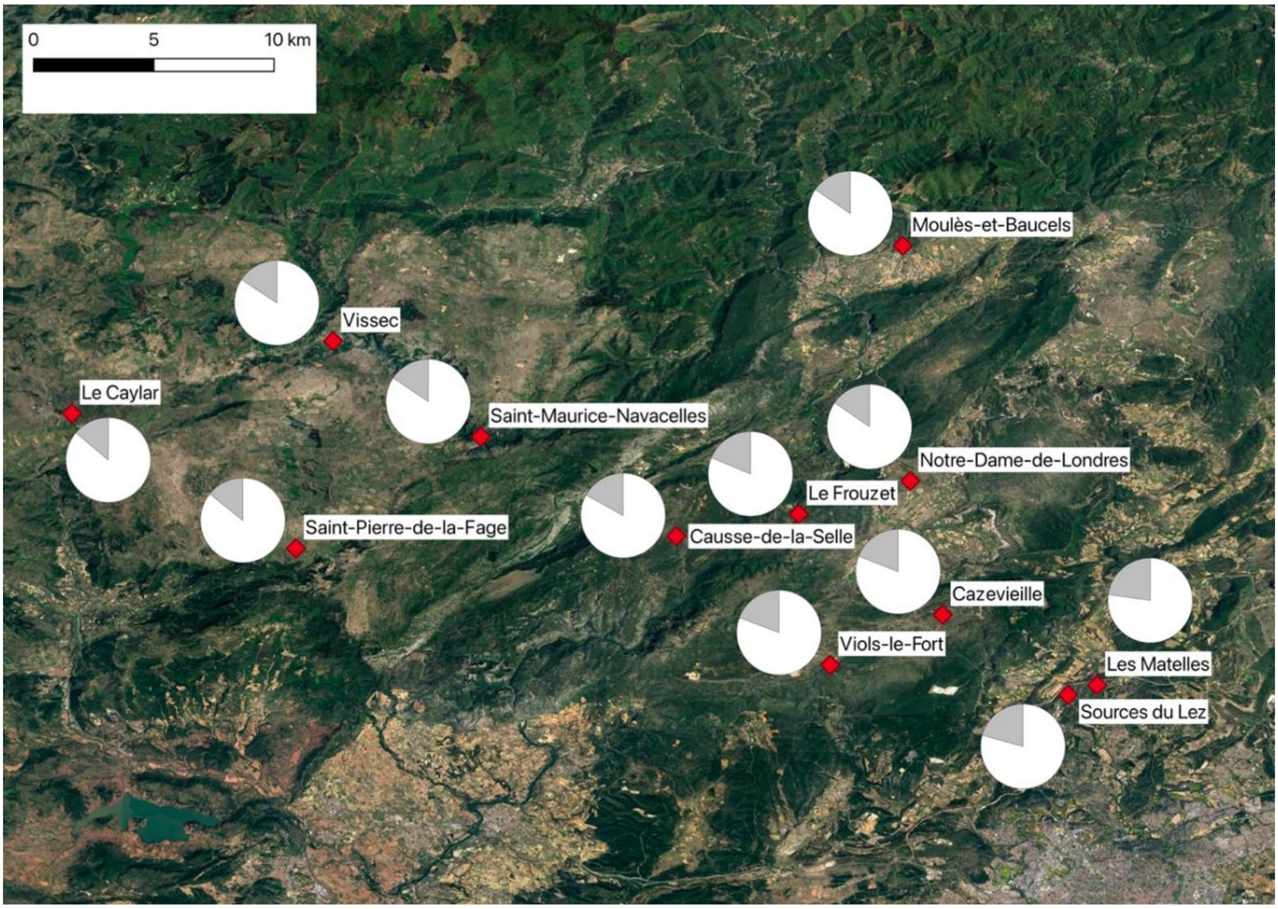


Fig. S1. The different sampling sites used for the survey with pheromone traps. The pie charts illustrate the frequency of the *typica* and *fasciata* morphs (in white) and *fusca* morph (in grey).

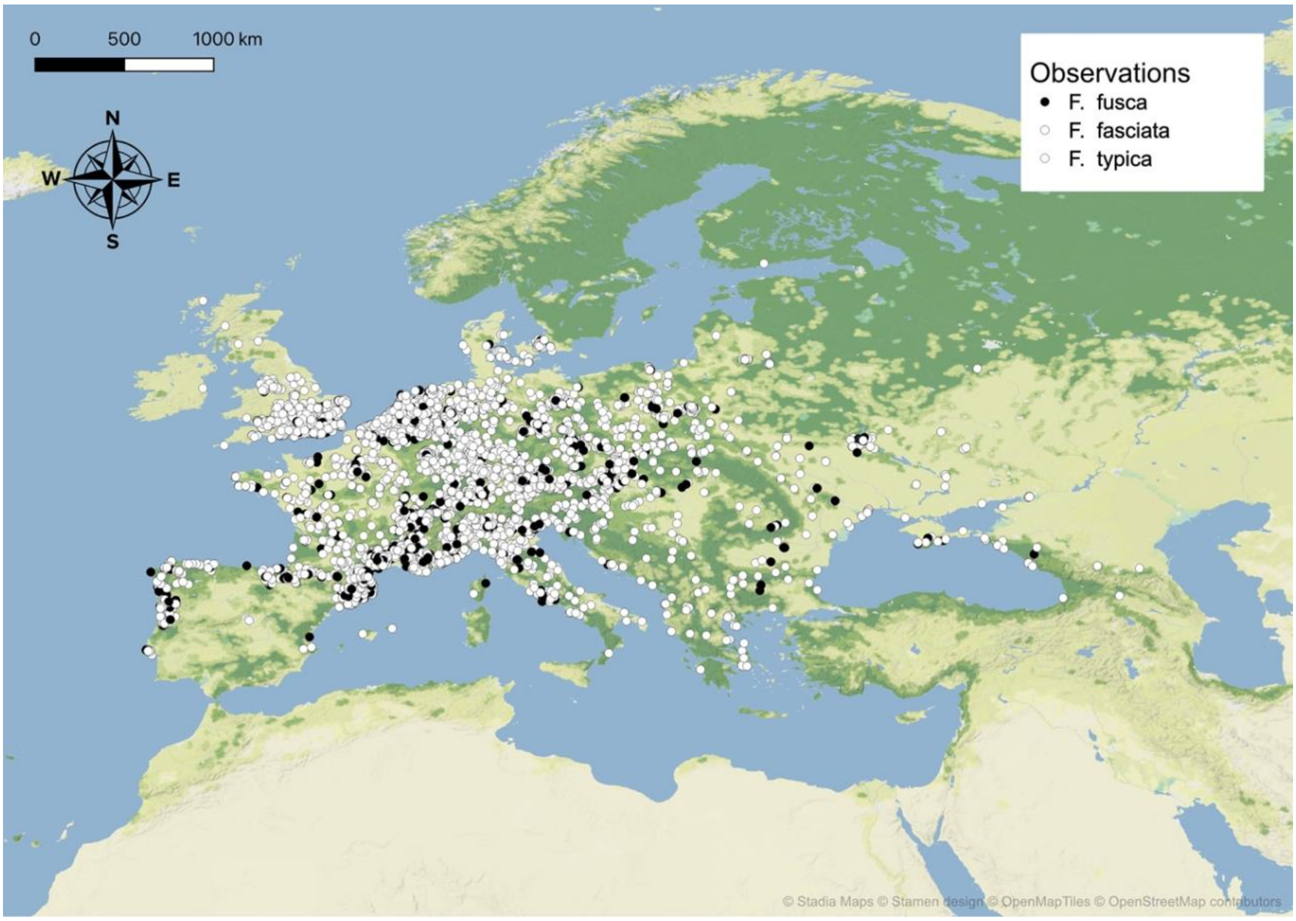


Fig. S2. The observations retrieved from citizen science.

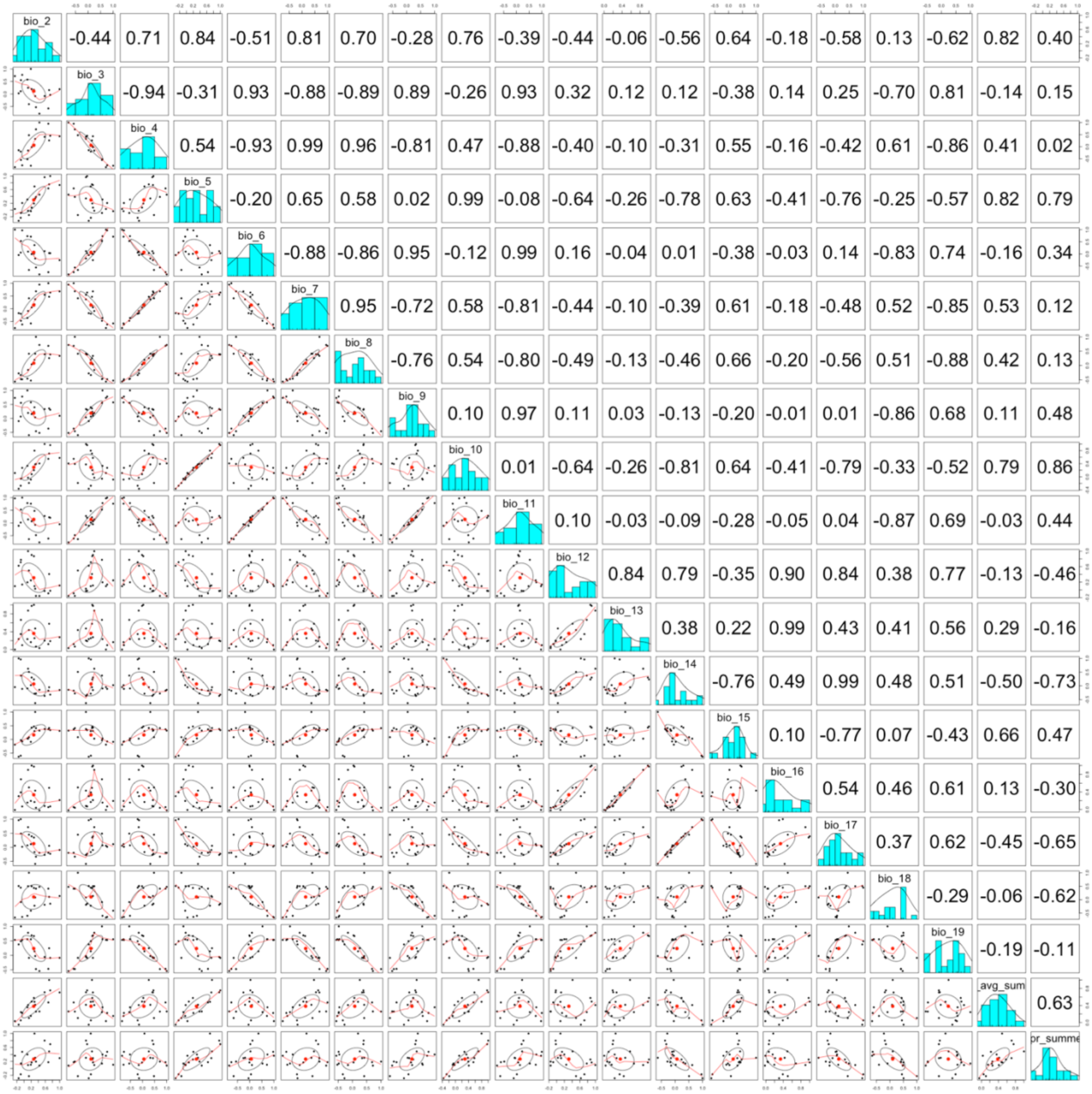


Fig. S3. Multicollinearity analysis results from variables for citizen science data. Below the diagonal: bivariate scatter plots, diagonal: histograms, above the diagonal: Pearson coefficients.

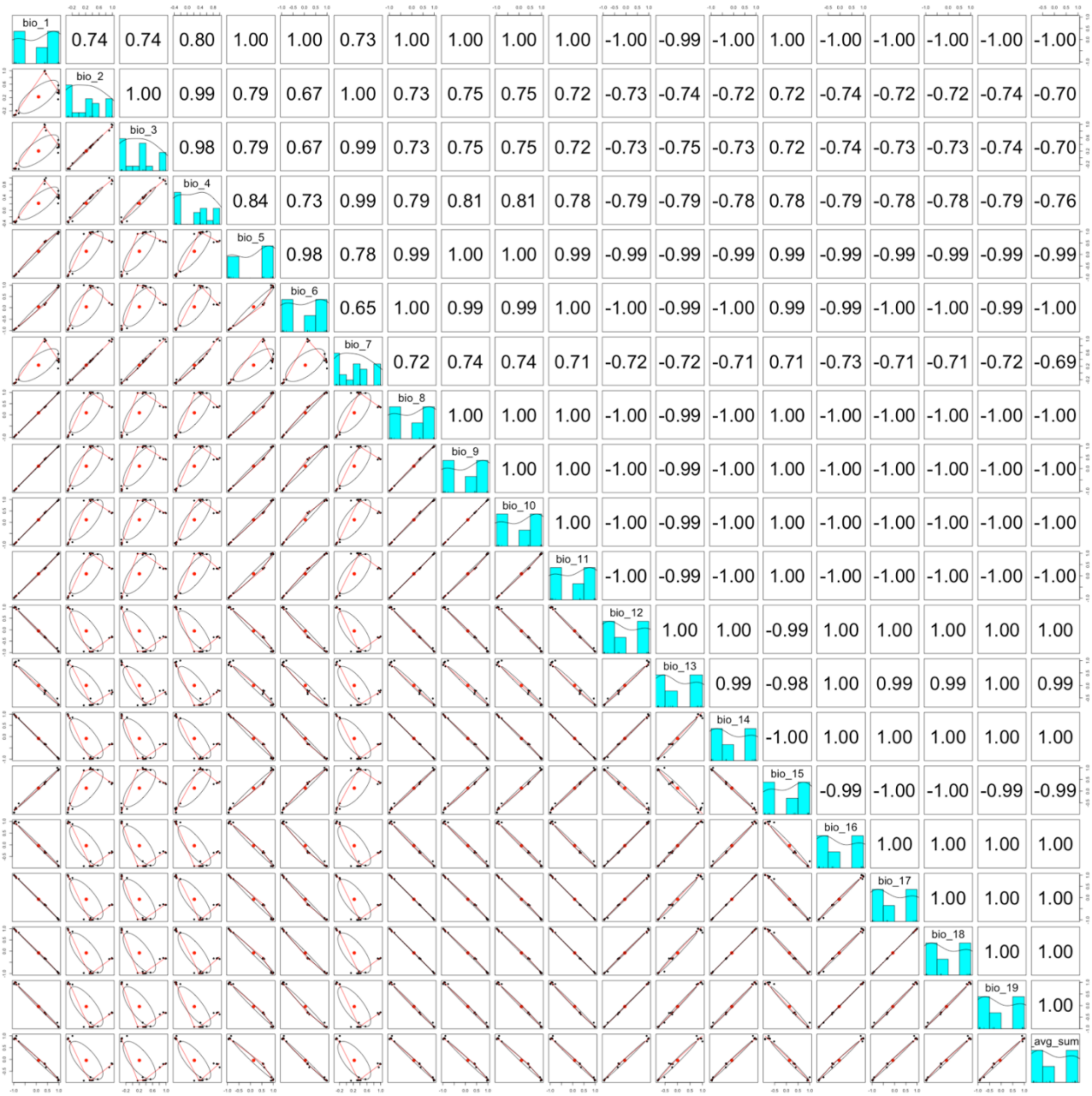


Fig. S4. Multicollinearity analysis results from variables for pheromone traps. Below the diagonal: bivariate scatter plots, diagonal: histograms, above the diagonal: Pearson coefficients.

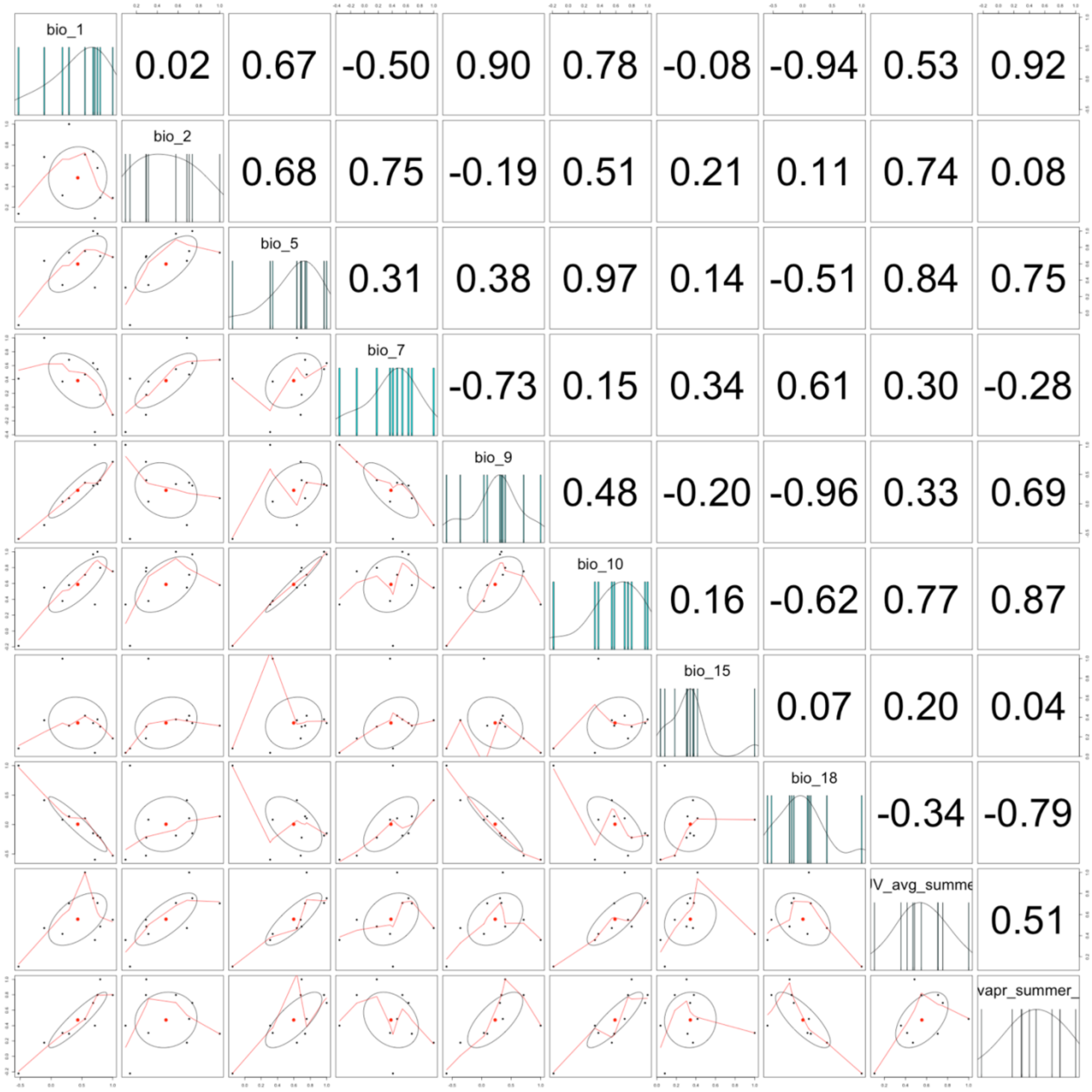


Fig. S5. Multicollinearity analysis results from variables for citizen science data, retained after preliminary model fitting. Below the diagonal: bivariate scatter plots, diagonal: histograms, above the diagonal: Pearson's coefficients.

| LOCALITY | LONG | LAT |
|---|---|---|
| Notre-Dame-de-Londres | 3.757654 | 43.829782 |
| Cazevieille | 3.774331 | 43.760857 |
| Le Frouzet | 3.700190 | 43.812721 |
| Viols-le-Fort | 3.716397 | 43.735085 |
| Sources du Lez | 3.838951 | 43.719664 |
| Saint-Pierre-de-la-Fage | 3.441096 | 43.795072 |
| Saint-Maurice-Navacelles | 3.536425 | 43.852600 |
| Vissec | 3.460274 | 43.901780 |
| Causse-de-la-Selle | 3.637097 | 43.801332 |
| Le Caylar | 3.325698 | 43.864505 |
| Moulès-et-Baucels | 3.753747 | 43.950984 |
| Les Matelles | 3.853955 | 43.724717 |

Table S1. Sites chosen for pheromone traps sampling and relative GPS coordinates.

| INTERROGATION, EUROPE |
|---|
| has[]=photos&quality_grade=any&identifications=any&place_id=96372&taxon_id=325295&verifiable=true&spam=false (Europe) |
| **INTERROGATION, ASIA** |
| quality_grade=any&identifications=any&swlat=23.07579611182776&swlng=88.70483423256081&nelat=43.12993913317876&nelng=172.4206545450608&taxon_id=325295&verifiable=true&spam=false (Asia). |

Table S2. Interrogations used on inaturalist site to retrieve data for the citizen science dataset.

| Factor | LR Chisq | Df | Pr(>Chisq) |
|---|---|---|---|
| **Pupal Weight (mg)** | 1.834 | 1 | 0.176 |
| **Morph** | 9.438 | 1 | 0.002 |
| **Sex** | 8.752 | 1 | 0.003 |

Table S3. Chi-Square Anova results for the GLM performed with lifespan as responsive variable.
